# An enhancer sequence in the intrinsically disordered region of the essential cell division protein FtsZ promotes conformation-guided substrate processing by ClpXP in *Escherichia coli*

**DOI:** 10.1101/2021.01.14.426688

**Authors:** Marissa G. Viola, Theodora Myrto Perdikari, Catherine Trebino, Negar Rahmani, Kaylee L. Mathews, Carolina Meija Pena, Xien Yu Chua, Botai Xuan, Christopher J. LaBreck, Nicolas L. Fawzi, Jodi L. Camberg

**Author notes:** **Corresponding author:** Jodi L. Camberg, 120 Flagg Road, Kingston, RI, 02881. These authors are co-first authors (Marissa G. Viola, Theodora Myrto Perdikari).

## Abstract

The essential bacterial division protein in *Escherichia coli*, FtsZ, assembles into the FtsZ-ring at midcell and recruits other proteins to the division site to promote septation. A region of the FtsZ amino acid sequence that links the conserved polymerization domain to a C-terminal protein interaction site was predicted to be intrinsically disordered and has been implicated in modulating spacing and architectural arrangements of FtsZ filaments. While the majority of cell division proteins that directly bind to FtsZ engage either the polymerization domain or the C-terminal interaction site, ClpX, the recognition and unfolding component of the bacterial ClpXP proteasome, has a secondary interaction with the predicted intrinsically disordered region (IDR) of FtsZ when FtsZ is polymerized. Here, we use NMR spectroscopy and reconstituted degradation reactions in vitro to demonstrate that this linker region is indeed disordered in solution and, further, that amino acids in the IDR of FtsZ enhance the degradation by conformationally-guided interactions.

## Introduction

The essential tubulin homolog in bacteria, FtsZ, assembles into head-to-tail protofilaments, and assembly is regulated by nucleotide binding, hydrolysis, and a network of regulator proteins that engage the FtsZ polymerization domain or an extended C-terminal region adjacent to the polymerization domain. This extended region has been a source of many investigations for insight into mechanisms to describe how bacterial cell division is regulated in response to FtsZ-interacting proteins, including an interaction with the actin homolog FtsA, which tethers the cytokinetic FtsZ-ring to the membrane. Several studies characterizing FtsZ from *Bacillus subtilis* and *Caulobacter crescentus* have shown that in addition to serving as a recognition site for FtsZ-interacting proteins, the extended C-terminal region may also have positional and sequence-specific effects on the arrangement and overall stability of FtsZ protofilaments in vitro and in vivo, which can further lead to filament bundling (1, 2).

The length and amino acid composition of the extended C-terminal region varies across organisms; however, it is widely predicted to be mostly an intrinsically disordered region (IDR) (3). In *Escherichia coli*, the length of the C-terminal region was reported to modify overall filament spacing in FtsZ protofilament bundles (4). Heterologous IDRs of similar length, but no sequence conservation, support division in *E. coli*, leading to a model by which the IDR serves the role of an entropic spring between the FtsZ protofilaments and the cytoplasmic membrane (3). While the C-terminal regions of different FtsZ proteins are widely predicted to include an IDR, there is significant variability among orthologs in the sequence identity of the region nearest the C-terminus. For example, the C-terminal peptide (CTP) region of FtsZ from *B. subtilis*, which includes the last 18 amino acids, contains positively charged residues that may autoregulate FtsZ polymerization and hydrolysis via electrostatic interactions (2). A docking site function conferred by the CTP region, however, is conserved across all FtsZ proteins and has species-specific protein interactions with C-terminal residues in FtsZ that modulate division via FtsZ binding (5-7). Moreover, several structures are available showing that in complex with other cell division proteins, the residues nearest the C-terminus, which participate in protein-protein interactions, likely exist as a helix or bent helix (8-11).

Several proteins bind to *E. coli* FtsZ via direct engagement of the FtsZ C-terminus, including but not limited to MinC, FtsA, ZipA, and ZapD; however, only one cell division modifier, ClpXP, has been implicated in binding directly to residues in the predicted IDR, as well as to residues near the C-terminus. ClpX from *E. coli* is a member of the AAA+ protein family (12-14). ClpX coassembles with the ClpP protease to recognize and degrade specific protein substrates, including FtsZ (15-17). During division, ClpXP degrades FtsZ from within the cytokinetic ring, which modifies the subunit exchange time of FtsZ in the highly dynamic ring structure (17), resulting in approximately 15% of total cellular FtsZ degraded by ClpXP per cell cycle (15-17).

Protein degradation is irreversible. To ensure high fidelity recognition, ClpX uses specific regions of a substrate, called degrons, to promote recognition. Degrons are typically present at either the N- or C-terminus of a substrate (18). FtsZ displays two motifs per polypeptide chain that are thought to engage ClpX, and FtsZ degradation is enhanced while polymerized, likely due to high local concentration and multivalency leading to higher avidity (16). One motif important for ClpXP degradation is near the C-terminus and overlapping with the multiprotein interaction site, and the second is in the predicted IDR linker region. Several proteins identified as ClpXP substrates have also been reported to contain IDRs, including RseA, IscU, and UmuD (18-23). Therefore, IDR-containing proteins may represent a distinct subset of ClpXP-controlled proteins. In this study, we elucidate the disordered structure of the full FtsZ C-terminal region from *E. coli* by solution NMR spectroscopy and dissect the multivalent complex interactions that regulate ClpXP degradation of polymerized and non-polymerized FtsZ. In this example, the IDR functions collaboratively with the C-terminal degron to precisely regulate degradation by ClpXP and promote conformation-guided substrate processing.

## Results

### NMR of the monomeric FtsZ C-terminus demonstrates that it is intrinsically disordered

The FtsZ C-terminal region (FtsZ CTR) is a 67 amino acid, 7.7 kDa FtsZ fragment, and is widely predicted to be intrinsically disordered (5). *E. coli* FtsZ CTR, which includes residues 317 through 383, has a flexible unstructured glutamine and proline rich linker and a highly conserved C-terminal core, also called the ‘CTP’ (C-terminal peptide) (370 through 379) that is essential for interactions with *E. coli* cell division proteins FtsA, ZipA, SlmA, MinC and ZapD (24). Although FtsZ CTR is dispensable for polymerization and is not required for the GTPase activity of FtsZ (25-27), it is important for recognition by cell division proteins and the ATP-dependent chaperone ClpX, which partners with ClpP to degrade FtsZ. A previous crystal structure of the C-terminal region of ZipA (residues 185 through 328) bound to the extreme C-terminus of *E. coli* FtsZ (residues 367 through 383) revealed that the 17-residue fragment of FtsZ binds ZipA as an extended β-strand at region 367-373 followed by an α-helix formed by residues 374-383 (11). On the other hand, the crystal structure of the DNA-activated FtsZ-ring inhibitor SlmA, in complex with an FtsZ C-terminal peptide fragment, revealed that FtsZ adopts an extended conformation (9). The structure of the FtsZ C-terminal peptide (336-351) from *Thermotoga maritima* in complex with FtsA revealed mostly helical content, with two small helical regions juxtaposed with a large bend in between (10). And finally, a recent NMR study on the FtsZ-ring positioning promoter protein MapZ, isolated from *Streptococcus pneumoniae*, showed that MapZ interacts with FtsZ C-terminal region regardless of its polymerization status but the structural details of this interaction were not investigated (28). In summary, few structures of FtsZ CTP bound to accessory proteins are available, and some of them depict the CTP bound as a helix while others report an extended conformation, suggesting that the FtsZ short core region adjacent to the C-terminus samples various conformational modes depending on its binding partner. However, currently there is no direct structural information on monomeric (unbound) FtsZ C-terminal sequences available.

Here, we used solution NMR spectroscopy to elucidate the secondary structure of the monomeric FtsZ CTR. The two-dimensional ^1^H-^15^N (heteronuclear single-quantum coherence – HSQC) NMR correlation spectra of FtsZ CTR exhibits narrow chemical shift dispersion, typical of a disordered protein, suggesting a lack of structural order consistent with sequence-based predictions (Fig. 1A). Furthermore, we assigned the ^13^C_α_ and ^13^C_β_ chemical shifts for FtsZ CTR and computed the secondary chemical shifts (ΔδC_α_ – ΔδC_β_), obtained by measuring the difference in the observed ^13^C_α_ and ^13^C_β_ chemical shifts and those predicted for a completely disordered structure. These secondary chemical shifts are mostly near zero for all residues, consistent with intrinsic disorder, except A376 and F377 which have positive secondary chemical shift values of about 1.5 ppm consistent with some population of α-helices (29, 30) as values above ∼4.0 ppm are expected with persistent helical structure. (Fig. 1B). Interestingly, this short helical region is localized in the highly conserved C-terminal core and is adjacent to the multiprotein recognition site. A previous alanine scanning mutagenesis study showed that mutation of Phe 377 to Ala reduced ZipA binding affinity, which is consistent with the fact that the hydrophobic residue F377 is highly conserved and forms helix-stabilizing contacts in a structure of FtsZ 367-383 bound to ZipA 185-328 (11). To estimate the helical population, we used the δ2D algorithm (31), which takes as input the chemical shifts. Based on this approach, we find that although FtsZ has a region with some helical population, even this region is *primarily* disordered with only a minor population of helical structure (375-379) (Fig. 1C). Hence, the C-terminal region of FtsZ does not adopt a highly helical structure in the absence of binding partners, and ^1^H-^15^N HSQC spectral overlay of the CTR indicates that it does not self-interact (Fig. S1). Taken together, our data suggest that FtsZ CTR is monomeric and remains mostly intrinsically disordered, although we observe a short, partially helical motif, amino acids 375 through 379, near the multiprotein interaction site that overlaps the C-terminal ClpX recognition site (amino acids 375-383).

**Figure 1.**
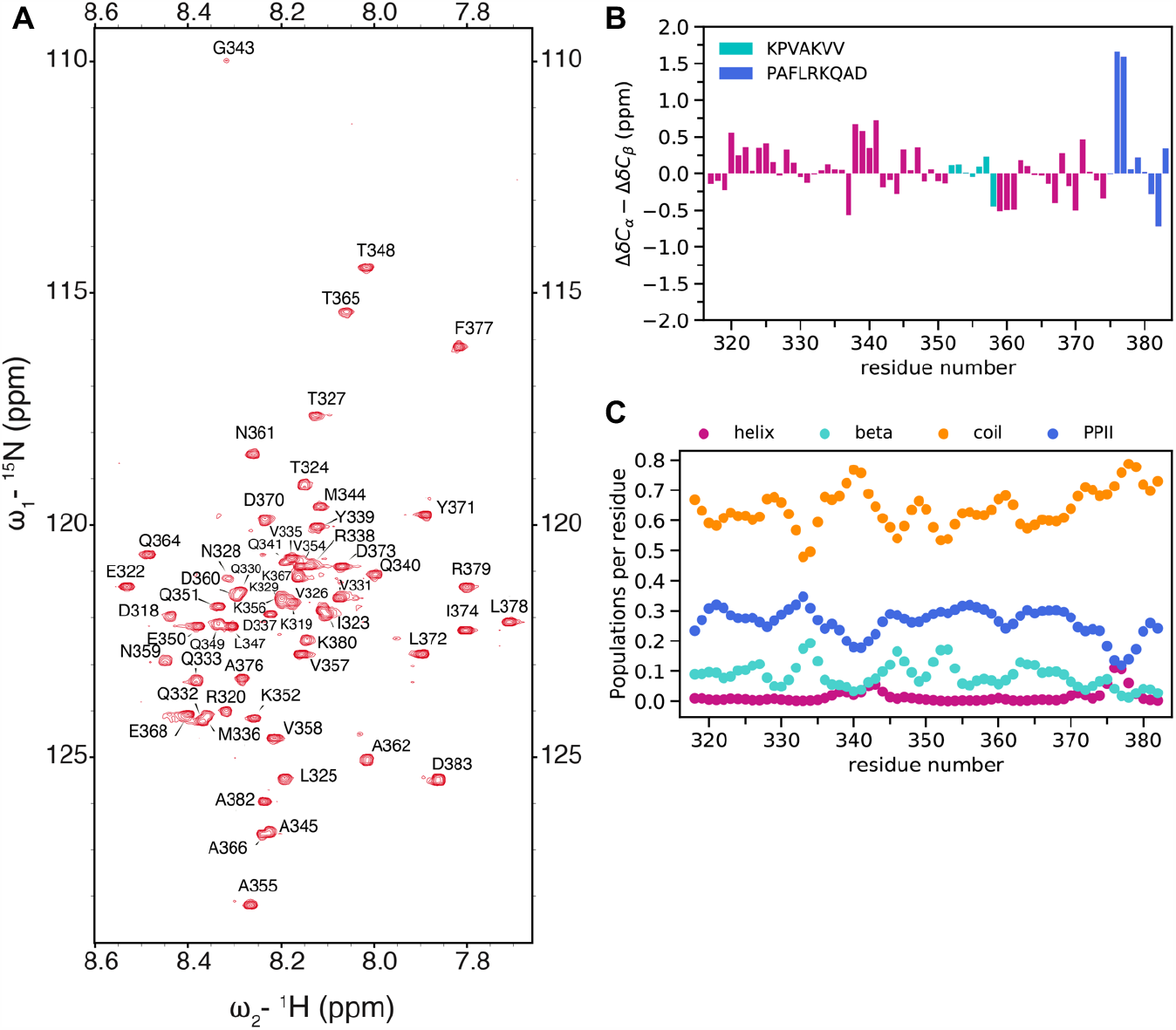
The monomeric FtsZ C-terminus is intrinsically disordered. (A) ^1^H-^15^N HSQC spectrum of FtsZ CTR is consistent with intrinsic disorder. (B) Secondary chemical shifts (ΔδC_α_ - ΔδC_β_) are localized around zero except A376 and F377 that have a much higher positive value consistent with local partial helical conformation. ^352^KPVAKVV^358^ sequence is highlighted in cyan and ^375^PAFLRKQAD^383^ sequence at the extreme C-terminus is highlighted in blue. (C) Secondary structure populations derived from the observed chemical shifts using the δ2D algorithm confirms the dominance of random coil (i.e., disordered) conformations.

### The intrinsically disordered C-terminal region of FtsZ is sufficient to target a model monomeric substrate for ClpXP degradation

To determine if the IDR of FtsZ is sufficient to target Gfp for degradation, and to further elucidate the relative contributions of both motifs implicated in ClpX recognition, which are separated by 16 amino acids, we utilized an engineered model substrate containing Gfp as a proxy for the FtsZ polymerization domain linked to the IDR of FtsZ (full CTR residues 317 through 383), called Gfp-IDR_FtsZ_ (Fig. 2A). To measure degradation of Gfp-IDR_FtsZ_ by ClpXP, we monitored the loss of fluorescence with time during incubation with ClpXP and ATP. We observed that Gfp-IDR_FtsZ_ fluorescence decreased with time indicating that ClpXP unfolds and degrades Gfp-IDR_FtsZ_; however, Gfp alone is not a substrate for ClpXP and maintains maximal fluorescence throughout the incubation period (Fig. 2B). These results indicate that the IDR of FtsZ is sufficient to promote recognition and degradation of Gfp by ClpXP.

**Figure 2.**
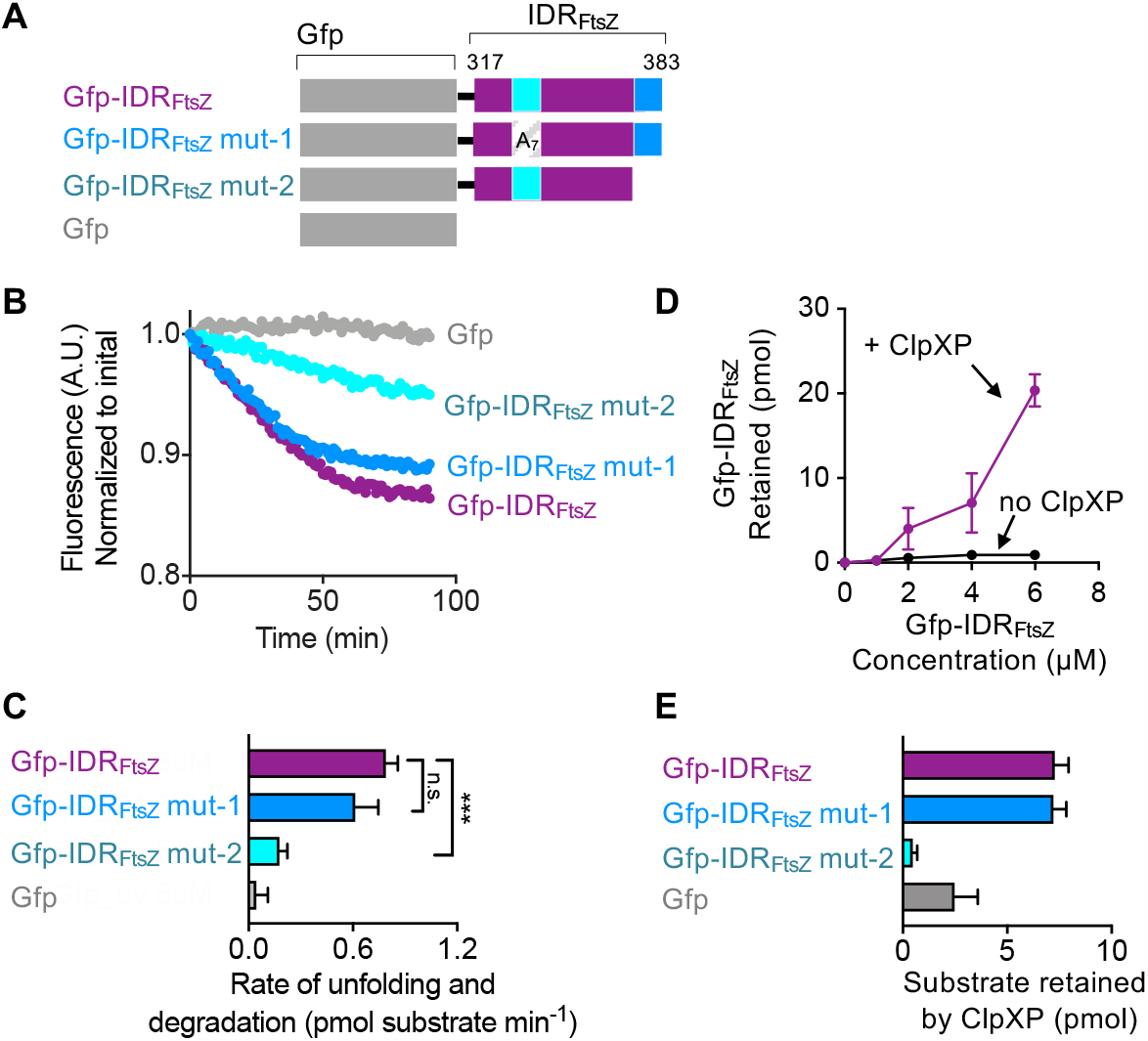
Degradation of a synthetic, monomeric FtsZ substrate by ClpXP. (A) Schematic for Gfp-IDR_FtsZ_ is shown. Gfp was fused to the C-terminal 67 amino acids of *E. coli* FtsZ IDR (residues 317-383) of FtsZ. Two variants were constructed, induing Gfp-IDR_FtsZ_ mut-1, which contains Ala substitutions at amino positions 352 through 358, corresponding to position in *E. coli* FtsZ, and Gfp-IDR_FtsZ_ mut-2, which is deleted for the last 9 C-terminal residues of FtsZ. (B) Gfp-IDR_FtsZ_ (8 μM) (purple), Gfp-IDR_FtsZ_ mut-1 (8 μM) (blue), Gfp-IDR_FtsZ_ mut-2 (8 μM) (aqua), and Gfp (8 μM) (gray) degradation was measured by loss of fluorescence with time in the presence of ClpX (1 μM), ClpP (1.2 μM), ATP (5 mM), and an acetate kinase regenerating system as described. Curves shown are representative of at least three replicates. (C) Rates of degradation were calculated as described in Materials and Methods. (D) Direct binding of Gfp-IDR_FtsZ_ to ClpXP was assayed by a filter retention assay. Retained Gfp-IDR_FtsZ_was measured by fluorescence. (E) Substrate retention by ClpXP was assayed as described in (C).

Next, to determine the relative contributions of each motif in Gfp-IDR_FtsZ_ implicated in ClpX recognition, which correspond to residues 379-383 and 352-358 of FtsZ, we constructed two additional chimeras, Gfp-IDR_FtsZ_ mut-1, which contains an intact C-terminus but has alanine substitution mutations in the upstream motif (352-358), and Gfp-IDR_FtsZ_ mut-2, with contains a deletion of the C-terminal motif (375-383) (Fig. 2A). We monitored Gfp fluorescence of both substrates in the presence of ClpXP and ATP to measure substrate degradation. We observed that ClpXP unfolds and degrades Gfp-IDR_FtsZ_ mut-1 to a similar extent as we previously observed with Gfp-IDR_FtsZ_, but Gfp-IDR_FtsZ_ mut-2 is degraded less efficiently (Fig. 2B). We calculated the rates of degradation and found that degradation of Gfp-IDR_FtsZ_ mut-2 was reduced by 77.4% compared to Gfp-IDR_FtsZ_, however, Gfp-IDR_FtsZ_ mut-1 was degraded at a rate within error of the Gfp-IDR_FtsZ_ rate of degradation (Fig. 2C). These results show that deletion of the C-terminal motif impairs degradation, but a multisite mutation in the upstream motif of Gfp-IDR_FtsZ_ has no significant effect on degradation, suggesting that ClpX primarily utilizes the FtsZ C-terminal end motif for recognition of an engineered monomeric substrate, and the secondary motif, located upstream in the IDR is dispensable and alone is not sufficient to robustly promote degradation.

Next, to confirm that reduced degradation correlates with reduced binding to ClpX, we used an ultrafiltration assay to collect substrate-bound ClpXP complexes. We incubated ClpXP with ATP at 0 °C to prevent degradation, assembled complexes containing ClpXP and Gfp-IDR_FtsZ_, and then collected and quantified complexes by ultrafiltration and fluorescence. We observed that ClpXP retains Gfp-IDR_FtsZ_ in a concentration-dependent manner, indicating the formation of enzyme-substrate complexes (Fig. 2D). Next, we compared all four engineered substrates in the ClpXP retention assay and observed that Gfp-IDR_FtsZ_ and Gfp-IDR_FtsZ_ mut-1 are efficiently retained by ClpXP; however, Gfp-IDR_FtsZ_ mut-2 and Gfp are poorly retained (Fig. 2E). These results suggest that the failure of ClpXP to degrade Gfp-IDR_FtsZ_ mut-2, compared to Gfp-IDR_FtsZ_ and Gfp-IDR_FtsZ_ mut-1, is likely due to poor recognition of the upstream motif. Moreover, the presence of the upstream motif appears to have no influence on ClpX recognizing the C-terminal motif, since Gfp-IDR_FtsZ_ and Gfp-IDR_FtsZ_ mut-1 are similarly degraded by ClpXP. Finally, to confirm that Gfp-IDR_FtsZ_ and Gfp-IDR_FtsZ_ mut-1, but not Gfp and Gfp-IDR_FtsZ_ mut-2, are recognized by ClpXP, we also measured the rate of ATP hydrolysis by ClpX with each engineered substrate to detect if substrate increases the rate of ClpXP ATP hydrolysis by allosteric activation, as has previously been reported with the ClpXP substrate Gfp-ssrA (32). We observed that the rate of ClpX ATP hydrolysis increased by approximately 20-40% for Gfp-IDR_FtsZ_ and Gfp-IDR_FtsZ_ mut-1; however, the rate of hydrolysis in the presence of Gfp-IDR_FtsZ_ mut-2 was not significantly different from ClpX without substrate (Table S1). These results are consistent with poor recognition of Gfp-IDR_FtsZ_ mut-2 by ClpX, relative to Gfp-IDR_FtsZ_ and Gfp-IDR_FtsZ_ mut-1.

### The secondary motif in the FtsZ IDR acts as a conformation-specific enhancer for degradation of FtsZ by ClpXP

As reported in previous studies, polymerized FtsZ is degraded by ClpXP more efficiently than non-polymerized FtsZ (15, 16). The upstream IDR motif does not appear to be involved in degradation of Gfp-IDR_FtsZ_, therefore, we hypothesized that it contributes to degradation of polymerized FtsZ. To test this, we used a full length Gfp-FtsZ fusion protein containing the complete wild type FtsZ sequence, including the polymerization domain and the full C-terminal region IDR, to quantitatively monitor ClpXP degradation by loss of fluorescence. Gfp-FtsZ was previously shown to assemble into protofilaments with GTP in vitro, similar to wild type FtsZ, hydrolyze GTP, and serve as a substrate for ClpXP degradation (17). We constructed mutations in the IDR of Gfp-FtsZ analogous to the mutations characterized with the engineered monomeric substrate Gfp-IDR_FtsZ_, including Gfp-FtsZ mut-1 and Gfp-FtsZ mut-2, which contain alanine substitutions in the upstream motif, corresponding to residues 352-358 of FtsZ, and deletion of the C-terminal recognition site, corresponding to residues 375-383 of FtsZ, respectively (Fig. 3A). We monitored degradation of Gfp-FtsZ, Gfp-FtsZ mut-1, and Gfp-FtsZ mut-2 in the presence of GTP, which promotes polymerization of Gfp-FtsZ, by measuring loss of fluorescence with time and calculated the rate of degradation of each substrate by ClpXP (Fig. 3B). We observed that Gfp-FtsZ is degraded by ClpXP in the presence of GTP, but the rate of Gfp-FtsZ mut-1 degradation is reduced by 48%. In the absence of GTP to promote polymerization, Gfp-FtsZ is degraded similarly as Gfp-FtsZ mut-1 (Fig. 3C), and half as efficiently as Gfp-FtsZ with GTP (Fig. 3B, 3C, and 3D). Consistent with our previous result with Gfp-IDR_FtsZ_, removal of the C-terminal motif severely impairs degradation by ClpXP (Fig. 2B, 3B, and 3C). These results show that the amino acid sequence of the upstream motif in the FtsZ IDR enhances degradation of polymerized Gfp-FtsZ, but does not serve as an effective ClpX targeting motif alone.

**Figure 3.**
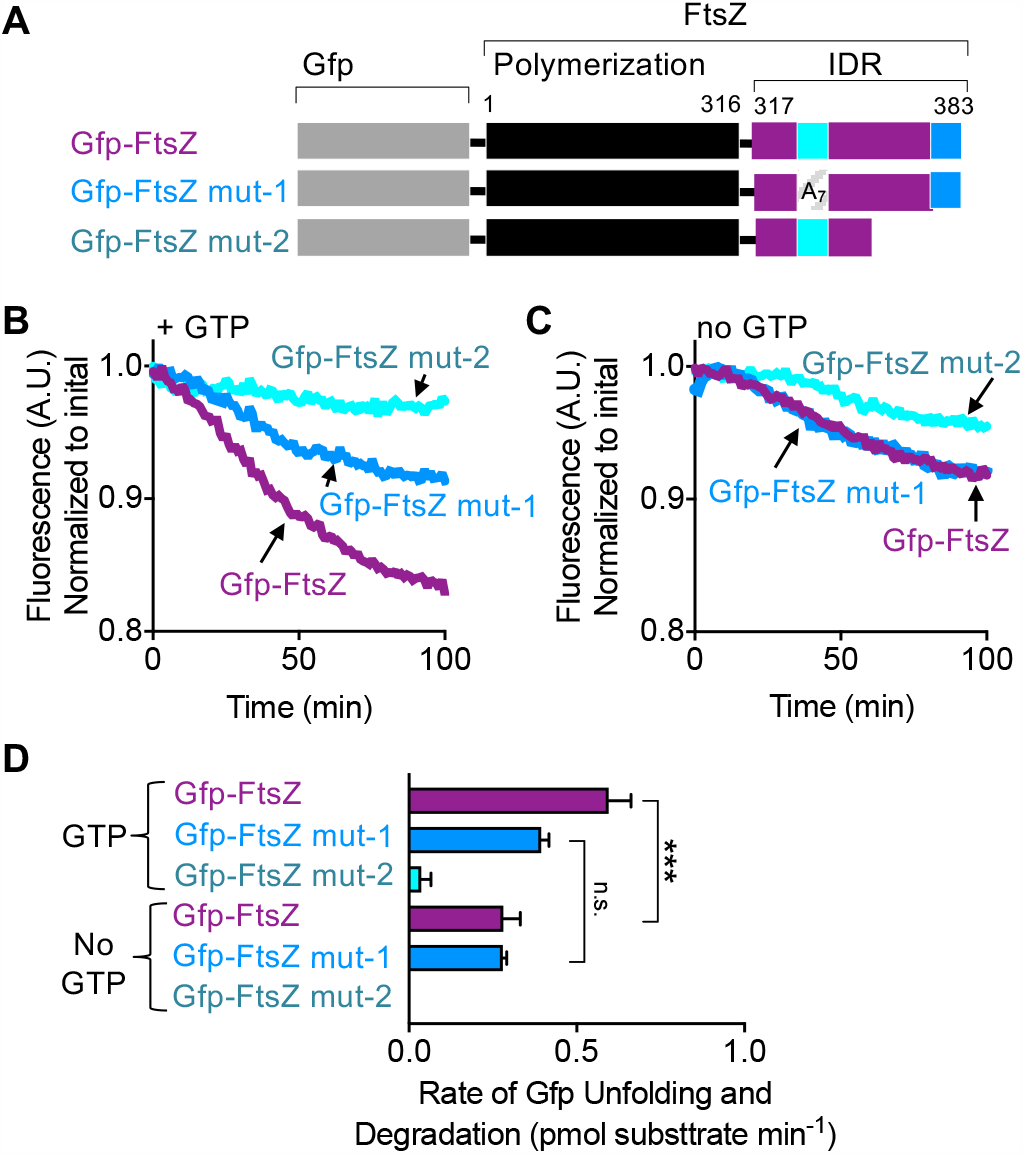
The motif in the IDR linker region is important for degradation of polymerized FtsZ by ClpXP. (A) Schematic for Gfp-FtsZ chimeric fusion proteins including Gfp-FtsZ, Gfp-FtsZ mut-1, which contains Ala substitutions at FtsZ positions 352 through 358, and Gfp-FtsZ mut-2, which is deleted for the last 9 C-terminal residues of FtsZ (B) with GTP and (C) without GTP. (D) Rates of unfolding and degradation were calculated as described in Materials and Methods. Curves shown are representative of at least three replicates.

### Architectural arrangement of recognition motifs in the IDR confers the conformational specificity that regulates degradation

The upstream and C-terminal ClpX recognition motifs in the FtsZ IDR are separated by 16 amino acids. To determine if the spacing between the two motifs in the IDR is important for degradation by ClpXP, we deleted residues 359 through 374 to construct Gfp-FtsZ Δ_space_ (Fig. 4A) and monitored degradation of Gfp-FtsZ Δ_space_ in reactions containing ClpXP and ATP. We observed that Gfp-FtsZ Δ_space_ is degraded by ClpXP at a similar rate as Gfp-FtsZ in both the absence and presence of GTP (Fig. 4A and 4B). These results suggest that the spacing between the recognition regions in the IDR is dispensable for the enhancement conferred by the presence of the secondary upstream motif. Finally, to test if there is a preference for relative position of one motif over another for degradation by ClpXP, we reversed the positions of both motifs to construct Gfp-FtsZ_swap_. We monitored degradation of Gfp-FtsZ_swap_ by ClpXP in the absence and presence of GTP (Fig. 4A and 4B) and observed that while Gfp-FtsZ_swap_ was degraded more efficiently than Gfp-FtsZ when GTP was omitted, there was no further enhancement observed when GTP was included in the reaction to promote polymerization. Together, these results suggest that residues 352 through 358 in the FtsZ IDR constitute a motif that enhances degradation of FtsZ; however, the enhancement is dependent on relative position within polymerized FtsZ.

**Figure 4.**
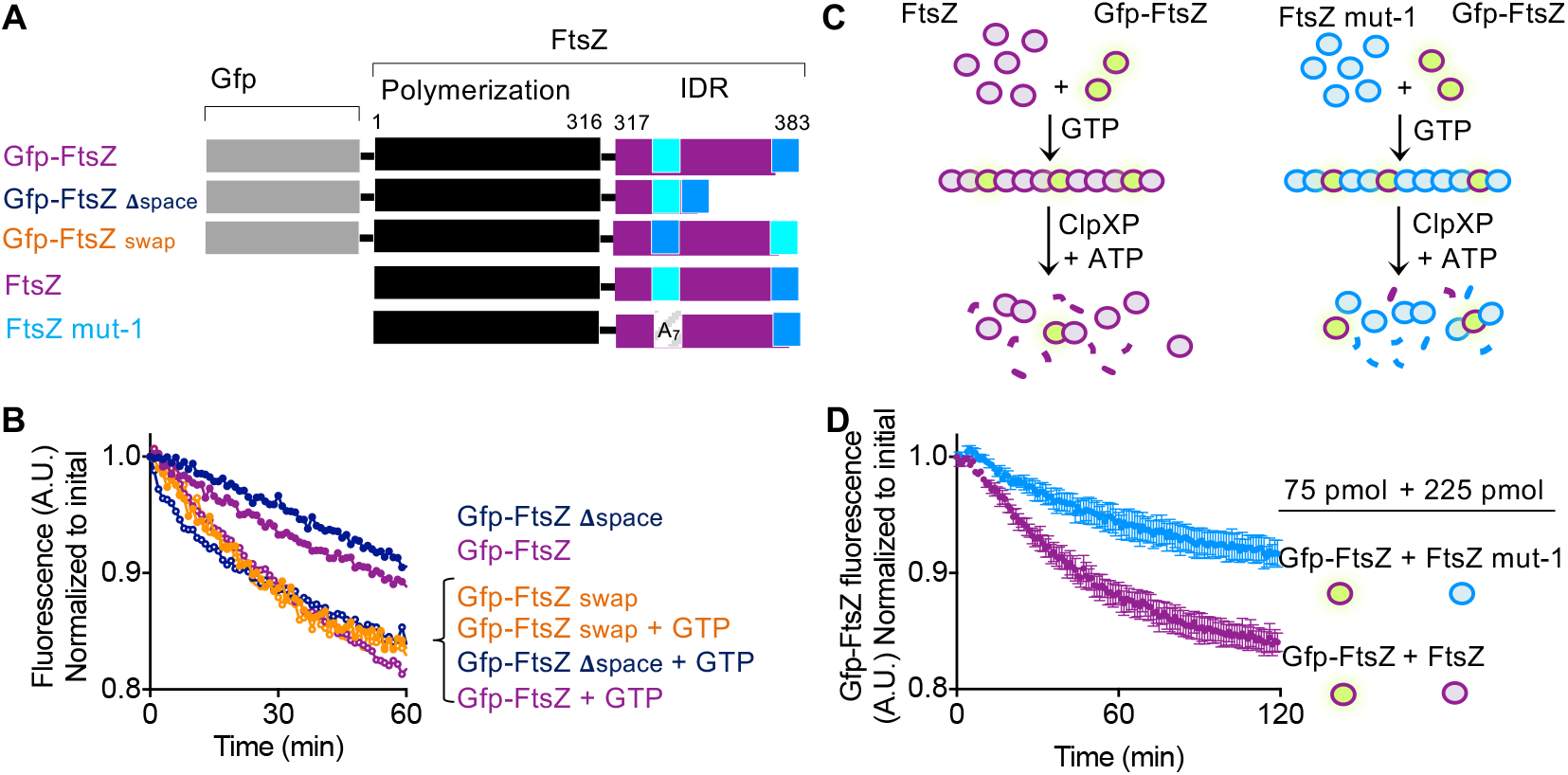
Relative degron position and distance affects FtsZ degradation by ClpXP. (A) Schematic for Gfp-FtsZ substrate is shown. Fluorescent protein Gfp is fused to the full-length FtsZ protein, which includes the polymerization domain (residues 1-316) and the IDR (residues 317-383). The 16 amino acid space between the two ClpX recognition motifs (‘352-358’ and ‘375-383’ as numbered in *E. coli*) in Gfp-FtsZ was deleted to construct Gfp-FtsZ_Δspace_, and relative positions of the degrons were exchanged to construct Gfp-FtsZ_swap_. (B) Gfp-FtsZ (purple), Gfp-FtsZ_Δspace_ (navy), and Gfp-FtsZ_swap_ (orange) (6 μM) (white triangles) were pre-incubated in the absence or presence of GTP to induce polymerization, then ClpXP (1 μM, 1.2 μM ClpP) with ATP (5 mM) were added to the reactions and fluorescence was monitored with time as described. Curves shown are representative of at least three replicates. (C) Cartoon depicting assembly of mixed polymers with GTP and subsequent degradation by ClpXP. Two populations of mixed polymers were assembled in the presence of GTP; one containing 25% Gfp-FtsZ in the assembly reaction with 75% wild type FtsZ, and another containing 25% Gfp-FtsZ with 75% FtsZ mut-1, which contains Ala substitutions at residues 352 through 358. (D) Mixtures of Gfp-FtsZ with wild type FtsZ subunits (purple) and Gfp-FtsZ with FtsZ mut-1 subunits, as described in (C), were pre-incubated with GTP to induce polymerization, then ClpX (1 μM), ClpP (1.2 μM) with ATP (5 mM) were added to the reactions and fluorescence was monitored with time as described. Data shown is the average of four independent replicates. Error is shown as standard deviation.

Finally, if the upstream motif in the IDR acts as a sequence-specific enhancer for degradation of polymerized FtsZ, then it could be functioning in one of two ways within the FtsZ polymer: (1) enhancement could occur as a result of nearby ClpX-recruitment sites in adjacent FtsZ subunits, or (2) the conformation of the IDR changes in response to polymerization through a direct interaction with another IDR or polymerization domain. To test if *trans* subunit targeting occurs within a single polymer, i.e., an interaction with one subunit increases the likelihood of an interaction with the adjacent subunit, we pre-assembled mixed polymers containing Gfp-FtsZ (75 pmol) with either wild type FtsZ (225 pmol) or FtsZ mut-1 (225 pmol), which has Ala substitutions in IDR upstream motif (352-358), in an assembly reaction with GTP (Fig. 4C). With this stoichiometry, both types of mixed polymers would contain 25% Gfp-FtsZ subunits that we could monitor by fluorescence, but the other 75% would be comprised of either wild type FtsZ or FtsZ mut-1, neither carrying a fluorescent label. Moreover, it was important to keep the total concentration of FtsZ wild type and mutant protein fixed (at 6 μM total), since the rate of FtsZ degradation increases with total FtsZ concentration (15). Finally, we know that Gfp-IDR_FtsZ_ and Gfp-IDR_FtsZ_ mut-1 are recognized and degraded by ClpXP similarly (Fig. 2B and 2E), so it is unlikely that differences would arise as a result of competition. We incubated the mixed polymers with ClpXP and ATP and measured Gfp-FtsZ degradation by monitoring the loss of fluorescence with time. We observed that the rate of Gfp-FtsZ degradation changed in response to whether FtsZ or FtsZ mut-1 were present in the reaction (Fig. 4C and 4D). Gfp-FtsZ was degraded from the polymers containing 75% wild type FtsZ subunits approximately 2-fold faster than from the polymers containing FtsZ mut-1, which lacks the upstream IDR sequence. These results show that degradation of Gfp-FtsZ is enhanced when non-fluorescent wild type FtsZ subunits are present, but not when the subunits contain alanine substitution mutations at residues 352 through 358. These results are consistent with the upstream motif in the IDR acting as an enhancer for degradation likely by improved targeting of neighboring subunits within a single polymer. Finally, to exclude the possibility that the IDR binds directly to another FtsZ subunit in a polymer with GTP, which could change the conformation of the IDR and modify ClpX recognition, we tested if Gfp-IDR_FtsZ_, which does not have a polymerization domain, can bind to FtsZ polymers in a direct recruitment assay. Therefore, we incubated FtsZ (8 μM) with GTP to promote polymerization in the presence of Gfp-IDR_Ftsz_, and then collected polymers by ultracentrifugation. We did not detect Gfp-IDR_Ftsz_ (8 μM) copelleting with FtsZ polymers, suggesting there is no interaction (Fig. S2). Additionally, we also tested if truncated FtsZ (FtsZΔC67) (8 μM), which does not contain its own IDR but polymerizes with GTP, can recruit Gfp-IDR_Ftsz_ in an ultracentrifugation assay, but also did not detect Gfp-IDR_Ftsz_ copelleting with FtsZΔC67 polymers (Fig. S2). These results suggest that the IDR of *E. coli* FtsZ does not engage other FtsZ subunits in a polymer under the conditions tested. Moreover, no IDR self-interaction was detected previously by solution NMR (Fig. S1).

### Although the FtsZ IDR binds other cell division proteins, it does not localize to the FtsZ-ring in the absence of the polymerization domain

In addition to ClpXP, other cell division proteins, such as FtsA, engage FtsZ via the multiprotein interaction site at the FtsZ C-terminus. During early cell division, FtsZ polymers assemble at midcell and form the FtsZ-ring (26, 33). When expressed in vivo, Gfp-FtsZ localizes to midcell and is tethered to the inner face of the cytoplasmic membrane through protein interactions with FtsA and ZipA. To determine if the IDR of FtsZ is sufficient to localize to midcell, since the multiprotein interaction site, which includes the FtsA interacting region, near the C-terminus remains intact, we expressed Gfp-IDR_FtsZ_ from a plasmid in *E. coli* MG1655 and visualized live dividing cells by fluorescence microscopy. As a control, we also expressed Gfp-FtsZ to localize the FtsZ-rings in dividing cells. We observed that while Gfp-FtsZ robustly localizes to fluorescent FtsZ-rings at visible septal regions, Gfp-IDR_FtsZ_ localizes uniformly throughout the cytoplasm (Fig. 5A and 5B). These results suggest that the IDR of *E. coli* FtsZ does not localize to a FtsZ-ring in the absence of the FtsZ polymerization domain, even though it retains the multiprotein interaction site near the C-terminus.

**Figure 5.**
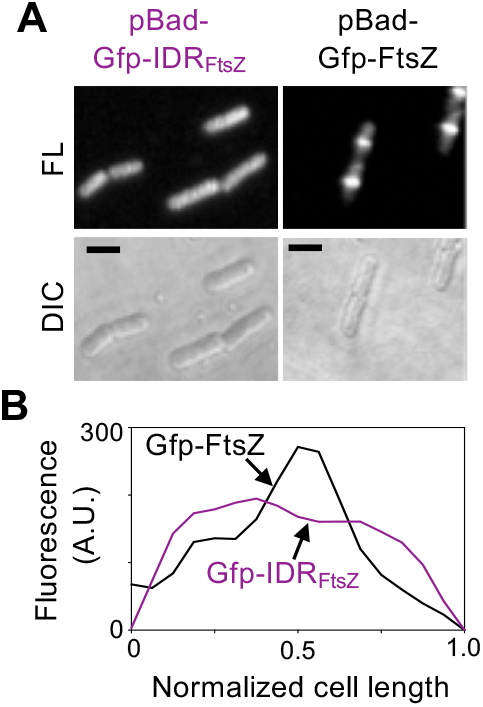
Polymerization is important for recruitment of the FtsZ IDR to FtsZ-rings in vivo. (A) Fluorescence (FL) and DIC microscopy images of *E. coli* MG1655 cells in log phase expressing Gfp-IDR_FtsZ_ and Gfp-FtsZ from an arabinose inducible plasmid show FtsZ-rings in cells expressing Gfp-FtsZ but not Gfp-IDR_FtsZ_. Size bars are 2 μm. (B) Cell fluorescence was measured and plotted as a function of cell length across the longitudinal axis of a cell expressing either Gfp-IDR_FtsZ_ (purple) or Gfp-FtsZ (black). FtsZ-rings are identified as a peak of fluorescence at 0.5 cell lengths. Traces shown are representative of at least three replicates.

## Discussion

Polymerization of FtsZ is a key feature of its function in vivo; however, the region of FtsZ that lies outside of the polymerization domain also serves important roles. These roles include mediating direct interactions with other proteins comprising the septal machinery, including regulators of FtsZ location and polymerization, functioning as a spacer between the membrane-tethering region and the main polymer filament, and, in several organisms, promoting lateral bundling through electrostatic interactions. Here, we show that the uncomplexed CTR of *E. coli* FtsZ is intrinsically disordered by solution NMR, as has previously been predicted (3), and that there is a small region of modest helical propensity near the terminus (Fig. 1A, 1B, and 1C).

While overall length of the IDR is important for division in *E. coli* (3), one protein, ClpX, has been implicated in engaging a central region within the IDR (16). We show that amino acids 352 through 358 of FtsZ contain sequence specific information that enhances degradation of FtsZ by ClpXP, but only when FtsZ is polymerized with GTP. The region near the FtsZ C-terminus thus functions as the major recognition element that promotes degradation of FtsZ by ClpXP. In our model, as ClpXP engages an FtsZ subunit, adjacent subunits in a polymer likely make contacts between the enhancer sequence of FtsZ, amino acids 352 through 358, and ClpX (Fig. 6). This substrate tethering event would prevent the second FtsZ subunit from escaping while ClpXP processes the first engaged FtsZ subunit. Processing, which includes processive unfolding and degradation, of polymerized FtsZ subunits by ClpXP would destabilize nearby head-to-tail FtsZ interactions, thus severing the polymer. This is consistent with the rapid and efficient FtsZ polymer destabilization activity that was previously observed and reported for ClpXP (17). While degradation of FtsZ by ClpXP is not essential for division to occur in *E. coli*, deletion of *clpX* or *clpP* from strains deleted for *minC* leads to filamentation and impaired division, indicating that ATP-dependent proteolysis regulates the division process (34). The *minC* gene encodes a component of the Min system, MinC, which destabilizes FtsZ polymers near the cell poles and ensures that the FtsZ-ring assembles at midcell. Thus, ClpXP has evolved a highly specialized mechanism to ensure that polymerized FtsZ is more effectively degraded than non-polymerized FtsZ.

**Figure 6.**
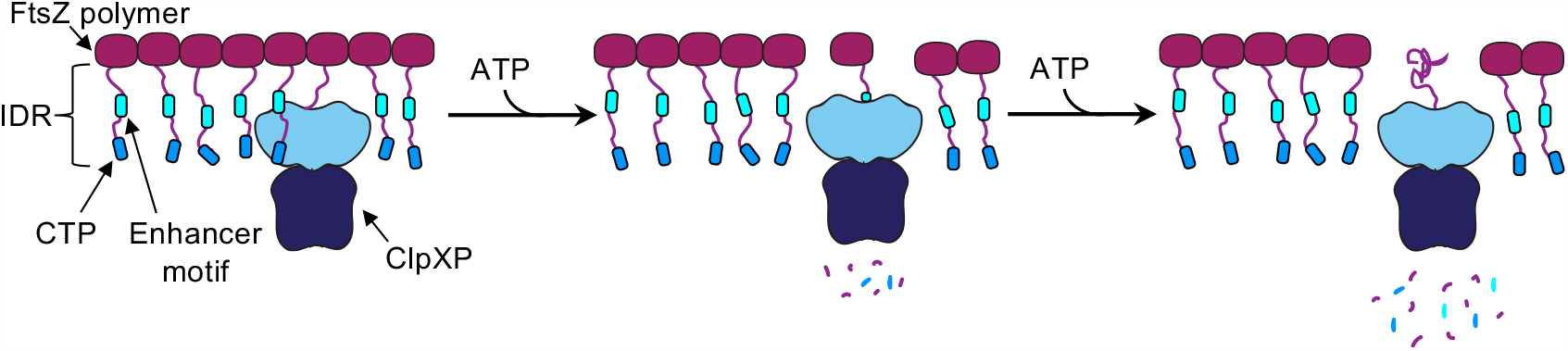
Model for conformation-guided recognition of FtsZ IDR by ClpXP. The primary degron that is responsible for recruiting ClpXP to FtsZ is present at the FtsZ C-terminus and includes residues 375 through 383, which overlaps with the broadly recognized CTP. When FtsZ is polymerized, a region of the IDR, referred to as the enhancer motif and including FtsZ residues 352 through 358, enhances recognition and degradation of FtsZ by ClpXP. In this model, ClpXP first engages a protomer and initiates unfolding and degradation. Then ClpXP engages a second, nearby subunit through direct recognition of the enhancer sequence on a neighboring protomer. This tethered adjacent protomer is prevented from release and also subsequently degraded. Polymerization of FtsZ leads to an accelerated rate of FtsZ protein turnover and FtsZ polymer severing catalyzed by ClpXP, thus releasing smaller fragments.

Among the pool of known ClpXP substrates in *E. coli*, several are reported to contain IDRs, including RseA, UmuD, IscU and FtsZ. It has been reported that IDR-containing proteins may inherently be sensitive to proteolysis in eukaryotes (35); however, protein half-lives in vivo do not strictly correlate with the presence of an IDR, and rather, recognition by proteases is likely context specific (36). Furthermore, length and position of IDRs could influence overall function and underly genetic variation (35). Other putative roles for IDRs include serving as an organizing hub. The IDR-hub model has been proposed for eukaryotic proteins, such as p53 and 14-3-3 (37, 38), IDR-mediated phase-separated condensates (39), and also for prokaryotic organizing proteins, such as PopZ. PopZ in *C. crescentus* is reported to network a highly dynamic, organized group of polar-localizing proteins via the IDR (40). While the FtsZ-ring recruits numerous proteins to the C-terminus of the IDR-containing region of FtsZ and forms phase-separated condensates in vitro (41), it is unclear if the condensates are relevant in vivo. Moreover, in the absence of the polymerization domain, the FtsZ IDR fails to self-associate or to localize to specific regions of the cell (Fig. 5). Polymerization of FtsZ concentrates FtsZ-interaction sites, such as those at the C-terminus, converting FtsZ to a multivalent ligand binding surface (42). However, ClpXP utilizes additional sequence-specific contacts in the IDR to enhance degradation of polymerized FtsZ. In addition to degrading FtsZ and destabilizing FtsZ polymers, ClpXP also disassembles and degrades FtsZ aggregates after heat shock (43). It is therefore likely that ClpXP may have evolved in *E. coli* to be a disassembly factor for accumulated FtsZ, ensuring turnover of densely networked FtsZ subunits and release of FtsZ and FtsZ-interacting proteins.

## Materials and Methods

### Strains, plasmids and chimeric fusion proteins

Strains and plasmids used in this study are listed in Table S2. Gfp-IDR_FtsZ_ and Gfp-FtsZ were cloned into expression plasmids for protein purification as described (17). For NMR studies, a TEV cleavage site was inserted into the Gfp-IDR_FtsZ_ by PCR mutagenesis. Substitution mutations in the IDR were constructed by site-directed mutagenesis using the QuikChange II XL Site-Directed Mutagenesis Kit (Agilent). The pBad-Gfp-IDR_FtsZ_ was constructed by cloning the coding sequence of Gfp-IDR_FtsZ_into the *NheI* and *HindIII* sites on the arabinose inducible vector, pBad24 (44). The pBad-Gfp-FtsZ expression plasmid was constructed as described (34). All mutations were confirmed by direct sequencing.

### Protein expression and purification

Uniformly ^15^N,^13^C labeled His-tagged Gfp-IDR_FtsZ_ was overexpressed in BL21 Star (DE3) *E. coli* (Invitrogen) in M9 minimal medium in H_2_O with ^15^N ammonium chloride and ^13^C glucose as the sole nitrogen and carbon sources, respectively. Cell pellets were harvested from 1L cultures induced with 1 mM IPTG at an OD_600_ of 0.6-1 after 3 hours at 30°C. Gfp-IDR_FtsZ_ pellets were resuspended in 50 mM Tris-Cl pH 8.0, 100 mM NaCl, 10 mM Imidazole, 1 mM DTT with 1x Roche Complete EDTA-free protease inhibitor. Resuspended pellets were lysed on an Emulsiflex C3 and the cell lysate was cleared by centrifugation (47,850 *x g* for 50 min at 4°C). The cleared supernatant was filtered using a 0.2 μM filter and loaded onto a 5 ml HisTrap HP Ni-affinity column (GE Healthcare). Protein was eluted in a gradient of 10 to 300 mM imidazole over five column volumes. Fractions containing Gfp-IDR_FtsZ_ were pooled together on a 3000 molecular weight cut-off (MWCO) dialysis membrane, followed by the addition of His-tagged TEV protease to a final concentration of 30 μg/ml and dialyzed overnight at 4°C into 50 mM Tris-Cl (pH 8.0), 100 mM NaCl, 1 mM DTT. The cleavage products were subjected to subtraction purification using a 5 ml HisTrap HP (GE Healthcare) to yield untagged IDR_FtsZ_ in the flow through. Purified IDR_FtsZ_was further dialyzed overnight at 4°C into 50 mM MES (pH 6.5) using a 3000 MWCO dialysis membrane and concentrated using a 3000 MWCO spin concentrator. Concentrated IDR_FtsZ_ were flash-frozen in small aliquots for subsequent NMR experiments.

All Gfp-FtsZ, Gfp-IDR_FtsZ_, and engineered variants for biochemical assays were expressed in *E. coli* BL21 (λ de3) and purified as N-terminally tagged six histidine fusion proteins by immobilized metal affinity chromatography as described (17). Wild type FtsZ, ClpX, and ClpP were expressed and purified as described (15, 45, 46). Protein concentrations are reported as Gfp-IDR_FtsZ_ monomers, Gfp-FtsZ monomers, FtsZ monomers, ClpX hexamers, and ClpP tetradecamers. All amino acid residue numbering refers to the *E. coli* FtsZ amino acid sequence position.

### NMR Sample Preparation and NMR Spectroscopy

Monomeric ^15^N,^13^C-labeled FtsZ CTR was thawed and diluted in 50 mM MES pH 6.5, 100 mM KCl, 10 mM MgCl_2_, 10% D_2_O to a final concentration of ∼163 μ?. Sample concentration was estimated using the extinction coefficient, 2560 M^-1^ cm^-1^, calculated by Protein Calculator v3.4 (http://protcalc.sourceforge.net/). NMR experiments were recorded at 25°C using Bruker Avance III HD NMR spectrometer operating at 850 MHz _1_H frequency equipped with a Bruker TCl z-axis gradient cryogenic probe. Experimental sweep widths, acquisition times, and the number of transients were optimized for the necessary resolution, experiment time, and signal to noise ratio for each type of experiment. NMR spectra were processed with NMRPipe (47) and analyzed with NMRFAM-Sparky (48).

^1^H-^15^N HSQC spectra were acquired with 3072 and 512 total points in the ^1^H and ^15^N dimensions with acquisition times of 172 and 135 ms, sweep widths of 10.5 and 22.0 ppm, and centered at 4.7 and 119.0 ppm, respectively. Each ^1^H-^15^N HSQC was acquired with the standard Bruker pulse sequence hsqcetf3gpsi.

Backbone amide resonance assignments were obtained by acquiring triple resonance assignment experiments HNCO, HN(CA)CO, CBCA(CO)NH, and HNCACB using standard Bruker TopSpin pulse sequences hncogp3d, hncacogp3d, cbcaconhgp3d, and hncacbgp3d. Each three-dimensional experiment was acquired with sweep widths of 10.0 ppm in the ^1^H dimension, 20.0 ppm in the ^15^N dimension, and 6.5 (CO) or 56.0 (CACB) ppm in the ^13^C dimension, centered at 4.7, 119.0, and 173.0 or 41.0 ppm, respectively. HNCO and HN(CA)CO experiments were acquired with 2048, 100, and 40 total points in the ^1^H, ^15^N, and ^13^C dimensions, while CBCA(CO)NH and HNCACB experiments were acquired with 2048, 84, and 120 total points, respectively.

### Fluorescence-based Protein Degradation assays

To monitor degradation by loss of fluorescence, Gfp (8 μM), Gfp-IDR_FtsZ_ (8 μM), Gfp-IDR_FtsZ_ variants (mut-1 and mut-2) (8 μM), Gfp-FtsZ (5 μM), and Gfp-FtsZ variants (mut-1 and mut-2) (5 μM), were incubated with ClpX (1 μM), ClpP (1.2 μM ClpP) and ATP (5 mM) HEPES buffer (50 mM, pH 7.0) containing 150 mM KCl, 10 mM MgCl_2_, 0.005% Triton X-100, and, where indicated, GTP (2 mM), acetate kinase (25 μg ml^-1^) and acetyl phosphate (15 mM). Gfp(uv) has been reported to dimerize at high concentrations, therefore we used concentrations of Gfp-fusion proteins in all experiments well below the dimerization condition (49). In degradation assays containing Gfp-FtsZ variants and omitting GTP, the bifunctional ATP/GTP-regenerating system of acetate kinase and acetyl phosphate was substituted with the ATP-specific system of creatine kinase (60 μg ml^-1^) and phosphocreatine (5 mg ml^-1^). Degron distance, position and subunit targeting assays were carried out with Gfp-FtsZ Δ_space_ (6 μM), Gfp-FtsZ_swap_ (6 μM), or a mixture of 1:3, Gfp-FtsZ:FtsZ, where indicated. Fluorescence was monitored in an Agilent Eclipse Spectrofluorometer at an excitation wavelength of 395 nm and emission of 510 nm. The background signal from buffer was subtracted from each data set and then data was normalized to report the change in the fraction of total arbitrary fluorescence units after time zero. Curve fitting was performed on GraphPad Prism (version 8.4.1). Rates of Gfp unfolding and degradation were calculated using the linear portions of curve (10 min to 30 min) from at least three replicates.

### Direct Binding Assays

To evaluate direct binding of Gfp-IDR and variants with ClpX, reactions (50 μl) containing ClpX (1 μM), ClpP (1 μM), ATP (5 mM) and, as indicated, Gfp-IDR_FtsZ_ or Gfp-IDR_FtsZ_ variants (0, 1, 2, 4 or 6 μM) in HEPES (50 mM, pH 7.0) buffer with KCl (150 mM), MgCl_2_ (10 mM), Triton X-100 (0.005%), and BSA (50 μg ml^-1^). After assembly for 15 minutes, all reactions were incubated on ice for 15 minutes. Reactions were transferred to polyethersulfone filters with a molecular weight cut off of 100 kDa (Pall) and centrifuged at 21,000 × *g* for 20 minutes at 23 C to collect complexes. Retained complexes were collected and bound Gfp variants were quantified by fluorescence.

### ATP Hydrolysis Assays

ATP hydrolysis rates for ClpX (0.5 μM) in HEPES buffer (50 mM, pH 7.5), with KCl (150 mM), MgCl_2_ (20 mM), and 5 mM ATP in the absence and presence of Gfp-IDR_FtsZ_, Gfp-IDR_FtsZ_ mut-1 and Gfp-IDR_FtsZ_ mut-2 (all 8 μM) were determined by measuring the amount of free phosphate released with time (0, 5, 10 and 15 minutes) using Biomol Green (Enzo Life Sciences). Phosphate was quantified by comparison to a phosphate standard curve.

### Sedimentation Assays

To determine if the IDR of FtsZ interacts with FtsZ polymers, reaction mixtures (25 μl) containing FtsZ or FtsZΔC67 (8 μM), with or without Gfp-IDR_FtsZ_ (8 μM) were prepared in HEPES buffer (50 mM, pH 7.0) containing KCl (150 mM), and MgCl_2_ (10 mM). Where indicated, 2 mM GTP, acetate kinase (25 μg ml^−1^), and acetyl phosphate (15 mM) were added to induce polymerization. Reactions were incubated for 10 minutes at 23°C and then centrifuged at 129,000 *× g* for 30 min at 23°C. Supernatants and pellets were analyzed by SDS-PAGE and coomassie staining.

### Fluorescence Microscopy

*E. coli* MG1655 strain (JC0390) was transformed with pBad-Gfp-IDR_FtsZ_ or pBad-FtsZ, and overnight cultures were diluted and grown to logarithmic conditions and induced with arabinose (0.002% and 0.001%, respectively) as described (16, 17). Images were collected with a Zeiss LSM 700 confocal fluorescence microscope and images were captured on an AxioCam digital camera with ZEN 2012 software.

## Data availability

The NMR chemical shift assignments for FtsZ CTR (317-383) from this publication have been deposited to the BMRB database (https://bmrb.io/) and assigned the accession number BMRB: 50314

## Acknowledgements

We thank Janet Atoyan for sequencing and microscopy assistance, Ben Piraino, Josiah Morrison and Colby Ferreira for helpful edits. Microscopy and sequencing were performed at the Rhode Island Genomics and Sequencing Center, which is supported in part by the National Science Foundation (MRI Grant No. DBI-0215393 and EPSCoR Grant No. 0554548 & EPS-1004057), the US Department of Agriculture (Grant Nos. 2002-34438-12688, 2003-34438-13111, and 2008-34438-19246), and the University of Rhode Island. This research is based in part on data obtained at the Brown University Structural Biology Core Facility, which is supported by the Division of Biology and Medicine, Brown University.

## References

1. K. Sundararajan et al., The bacterial tubulin FtsZ requires its intrinsically disordered linker to direct robust cell wall construction. Nature communications 6, 7281 (2015).

2. M. C. Cohan, A. M. P. Eddelbuettel, P. A. Levin, R. V. Pappu, Dissecting the Functional Contributions of the Intrinsically Disordered C-terminal Tail of Bacillus subtilis FtsZ. Journal of molecular biology 432, 3205–3221 (2020).

3. K. A. Gardner, D. A. Moore, H. P. Erickson, The C-terminal linker of Escherichia coli FtsZ functions as an intrinsically disordered peptide. Molecular microbiology 89, 264–275. (2013).

4. S. Huecas et al., Self-Organization of FtsZ Polymers in Solution Reveals Spacer Role of the Disordered C-Terminal Tail. Biophysical journal 113, 1831–1844 (2017).

5. X. Ma, W. Margolin, Genetic and functional analyses of the conserved C-terminal core domain of Escherichia coli FtsZ. J. Bacteriol. 181, 7531–7544 (1999).

6. S. A. Haney et al., Genetic analysis of the Escherichia coli FtsZ.ZipA interaction in the yeast two-hybrid system. Characterization of FtsZ residues essential for the interactions with ZipA and with FtsA. J. Biol. Chem. 276, 11980–11987 (2001).

7. J. Conti, M. G. Viola, J. L. Camberg, FtsA reshapes membrane architecture and remodels the Z-ring in Escherichia coli. Molecular microbiology 107, 558–576 (2018).

8. M. A. Schumacher, K. H. Huang, W. Zeng, A. Janakiraman, Structure of the Z Ring-associated Protein, ZapD, Bound to the C-terminal Domain of the Tubulin-like Protein, FtsZ, Suggests Mechanism of Z Ring Stabilization through FtsZ Cross-linking. The Journal of biological chemistry 292, 3740–3750 (2017).

9. M. A. Schumacher, W. Zeng, Structures of the nucleoid occlusion protein SlmA bound to DNA and the C-terminal domain of the cytoskeletal protein FtsZ. Proceedings of the National Academy of Sciences of the United States of America 113, 4988–4993 (2016).

10. P. Szwedziak, Q. Wang, S. M. Freund, J. Lowe, FtsA forms actin-like protofilaments. The EMBO journal 31, 2249–2260 (2012).

11. L. Mosyak et al., The bacterial cell-division protein ZipA and its interaction with an FtsZ fragment revealed by X-ray crystallography. EMBO J. 19, 3179–3191 (2000).

12. R. T. Sauer, T. A. Baker, AAA+ proteases: ATP-fueled machines of protein destruction. Annual review of biochemistry 80, 587–612 (2011).

13. T. A. Baker, R. T. Sauer, ATP-dependent proteases of bacteria: recognition logic and operating principles. Trends Biochem. Sci. 31, 647–653 (2006).

14. J. Snider, G. Thibault, W. A. Houry, The AAA+ superfamily of functionally diverse proteins. Genome biology 9, 216 (2008).

15. J. L. Camberg, J. R. Hoskins, S. Wickner, ClpXP protease degrades the cytoskeletal protein, FtsZ, and modulates FtsZ polymer dynamics. Proceedings of the National Academy of Sciences of the United States of America 106, 10614–10619 (2009).

16. J. L. Camberg, M. G. Viola, L. Rea, J. R. Hoskins, S. Wickner, Location of dual sites in E. coli FtsZ important for degradation by ClpXP; one at the C-terminus and one in the disordered linker. PloS one 9, e94964 (2014).

17. M. G. Viola, C. J. LaBreck, J. Conti, J. L. Camberg, Proteolysis-Dependent Remodeling of the Tubulin Homolog FtsZ at the Division Septum in Escherichia coli. PloS one 12, e0170505 (2017).

18. J. M. Flynn, S. B. Neher, Y. I. Kim, R. T. Sauer, T. A. Baker, Proteomic discovery of cellular substrates of the ClpXP protease reveals five classes of ClpX-recognition signals. Mol. Cell 11, 671–683. (2003).

19. S. M. Simon, F. J. Sousa, R. Mohana-Borges, G. C. Walker, Regulation of Escherichia coli SOS mutagenesis by dimeric intrinsically disordered umuD gene products. Proceedings of the National Academy of Sciences of the United States of America 105, 1152–1157 (2008).

20. S. B. Neher, R. T. Sauer, T. A. Baker, Distinct peptide signals in the UmuD and UmuD’ subunits of UmuD/D’ mediate tethering and substrate processing by the ClpXP protease. Proc. Natl. Acad. Sci. U. S. A. 100, 13219–13224 (2003).

21. R. Chaba, I. L. Grigorova, J. M. Flynn, T. A. Baker, C. A. Gross, Design principles of the proteolytic cascade governing the sigmaE-mediated envelope stress response in Escherichia coli: keys to graded, buffered, and rapid signal transduction. Genes & development 21, 124–136 (2007).

22. R. Pancsa, F. Zsolyomi, P. Tompa, Co-Evolution of Intrinsically Disordered Proteins with Folded Partners Witnessed by Evolutionary Couplings. Int J Mol Sci 19 (2018).

23. S. Sato et al., Evidence for dynamic in vivo interconversion of the conformational states of IscU during iron-sulfur cluster biosynthesis. Molecular microbiology 10.1111/mmi.14646 (2020).

24. M. Wang, C. Fang, B. Ma, X. Luo, Z. Hou, Regulation of cytokinesis: FtsZ and its accessory proteins. Curr Genet 66, 43–49 (2020).

25. X. Ma, D. W. Ehrhardt, W. Margolin, Colocalization of cell division proteins FtsZ and FtsA to cytoskeletal structures in living Escherichia coli cells by using green fluorescent protein. Proceedings of the National Academy of Sciences of the United States of America 93, 12998–13003 (1996).

26. H. P. Erickson, D. E. Anderson, M. Osawa, FtsZ in bacterial cytokinesis: cytoskeleton and force generator all in one. Microbiol Mol Biol Rev 74, 504–528 (2010).

27. X. Wang, J. Lutkenhaus, FtsZ ring: the eubacterial division apparatus conserved in archaebacteria. Molecular microbiology 21, 313–319 (1996).

28. T. Hosek et al., Structural features of the interaction of MapZ with FtsZ and membranes in Streptococcus pneumoniae. Scientific reports 10, 4051 (2020).

29. D. S. Wishart, B. D. Sykes, The 13C chemical-shift index: a simple method for the identification of protein secondary structure using 13C chemical-shift data. J Biomol NMR 4, 171–180 (1994).

30. D. S. Wishart, B. D. Sykes, F. M. Richards, The chemical shift index: a fast and simple method for the assignment of protein secondary structure through NMR spectroscopy. Biochemistry 31, 1647–1651 (1992).

31. C. Camilloni, A. De Simone, W. F. Vranken, M. Vendruscolo, Determination of secondary structure populations in disordered states of proteins using nuclear magnetic resonance chemical shifts. Biochemistry 51, 2224–2231 (2012).

32. R. E. Burton, T. A. Baker, R. T. Sauer, Energy-dependent degradation: Linkage between ClpX-catalyzed nucleotide hydrolysis and protein-substrate processing. Protein science : a publication of the Protein Society 12, 893–902 (2003).

33. C. Ortiz, P. Natale, L. Cueto, M. Vicente, The keepers of the ring: regulators of FtsZ assembly. FEMS microbiology reviews 40, 57–67 (2016).

34. J. L. Camberg, J. R. Hoskins, S. Wickner, The interplay of ClpXP with the cell division machinery in Escherichia coli. J. Bacteriol. 193, 1911–1918 (2011).

35. R. van der Lee et al., Intrinsically disordered segments affect protein half-life in the cell and during evolution. Cell reports 8, 1832–1844 (2014).

36. M. J. Suskiewicz, J. L. Sussman, I. Silman, Y. Shaul, Context-dependent resistance to proteolysis of intrinsically disordered proteins. Protein science : a publication of the Protein Society 20, 1285–1297 (2011).

37. V. N. Uversky, C. J. Oldfield, A. K. Dunker, Intrinsically disordered proteins in human diseases: introducing the D2 concept. Annu Rev Biophys 37, 215–246 (2008).

38. C. J. Oldfield et al., Flexible nets: disorder and induced fit in the associations of p53 and 14-3-3 with their partners. BMC genomics 9 Suppl 1, S1 (2008).

39. M. C. Cohan, R. V. Pappu, Making the Case for Disordered Proteins and Biomolecular Condensates in Bacteria. Trends Biochem Sci 45, 668–680 (2020).

40. J. A. Holmes et al., Caulobacter PopZ forms an intrinsically disordered hub in organizing bacterial cell poles. Proceedings of the National Academy of Sciences of the United States of America 113, 12490–12495 (2016).

41. B. Monterroso et al., Bacterial FtsZ protein forms phase-separated condensates with its nucleoid-associated inhibitor SlmA. EMBO Rep 20 (2019).

42. S. Du, K. T. Park, J. Lutkenhaus, Oligomerization of FtsZ converts the FtsZ tail motif (conserved carboxy-terminal peptide) into a multivalent ligand with high avidity for partners ZipA and SlmA. Molecular microbiology 95, 173–188 (2015).

43. C. J. LaBreck, S. May, M. G. Viola, J. Conti, J. L. Camberg, The Protein Chaperone ClpX Targets Native and Non-native Aggregated Substrates for Remodeling, Disassembly, and Degradation with ClpP. Frontiers in molecular biosciences 4, 26 (2017).

44. L. M. Guzman, D. Belin, M. J. Carson, J. Beckwith, Tight regulation, modulation, and high-level expression by vectors containing the arabinose PBAD promoter. Journal of bacteriology 177, 4121–4130 (1995).

45. M. R. Maurizi, M. W. Thompson, S. K. Singh, S. H. Kim, Endopeptidase Clp: ATP-dependent Clp protease from Escherichia coli. Methods Enzymol. 244, 314–331. (1994).

46. R. Grimaud, M. Kessel, F. Beuron, A. C. Steven, M. R. Maurizi, Enzymatic and structural similarities between the Escherichia coli ATP-dependent proteases, ClpXP and ClpAP. J. Biol. Chem. 273, 12476–12481. (1998).

47. F. Delaglio et al., NMRPipe: a multidimensional spectral processing system based on UNIX pipes. J Biomol NMR 6, 277–293 (1995).

48. W. Lee, M. Tonelli, J. L. Markley, NMRFAM-SPARKY: enhanced software for biomolecular NMR spectroscopy. Bioinformatics 31, 1325–1327 (2015).

49. F. G. Prendergast, K. G. Mann, Chemical and physical properties of aequorin and the green fluorescent protein isolated from Aequorea forskalea. Biochemistry 17, 3448–3453 (1978).

