## Supporting information for "An enhancer sequence in the intrinsically disordered region of the essential cell division protein FtsZ promotes conformation-guided substrate processing by ClpXP in *Escherichia coli*"

*These authors are co-first authors

### Table S1. ClpX ATP hydrolysis in the presence of Gfp-IDR$_{FtsZ}$ variants

| Substrate added | ClpX ATPase Activity (min$^{-1}$ ± S.E.M.) | p-value[1] |
|---|---|---|
| none | 37.0 ± 2.0 | |
| Gfp-IDR$_{FtsZ}$ | 43.5 ± 2.0 | 0.0204 |
| Gfp-IDR$_{FtsZ}$ mut-1 | 50.7 ± 0.3 | 0.0002 |
| Gfp-IDR$_{FtsZ}$ mut-2 | 40.7 ± 1.6 | n.s. |
| Gfp | 34.1 ± 1.9 | n.s. |

[1]P-values are compared to ClpX ATP hydrolysis without substrate; n.s. refers to not significantly different from the rate of ClpX in the absence of substrate.

**Table S2.** ***E. coli*** **strains and plasmids used for constructions in this study**

| *E. coli* Strain or Plasmid | Relevant Genotype Description | Source, reference or construction |
|---|---|---|
| Strains | | |
| BL21 (λde3) | F- *ompT gal dcm lon hsdSB(rB- mB-)* λ(de3[*lacI lacUV5-T7 gene 1 ind1 sam7 nin5*]) | EMD Millipore |
| BL21 Star (DE3) | | ThermoFisher |
| MG1655 | *LAM-rph-1* | [1] |
| Plasmids | | |
| pEt-FtsZ | *kan* | [2] |
| pEt-ClpX | *kan* | [2] |
| pEt-ClpP | *kan* | [2] |
| pEt-FtsZ(ΔC67) | *kan* | [3] |
| pEt-$His_6$-Gfp | *kan, gfp* | [3] |
| pEt-$His_6$-Gfp-IDR | *kan, gfp-IDR (FtsZ 317-383)* | [4] |
| pEt-$His_6$-Gfp-TEV-IDR | *kan, gfp-IDR (TEV site)* | This study |
| pEt-$His_6$-Gfp-IDR mut-1 | *kan, gfp-IDR ($_{352\text{-}AAAAAAA\text{-}358}$)* | This study |
| pEt-$His_6$-Gfp-IDR mut-2 | *kan, gfp-IDR (ΔC9)* | This study |
| pEt-$His_6$-Gfp-FtsZ | *kan, gfp-ftsZ* | [4] |
| pEt-$His_6$-Gfp-FtsZ mut-1 | *kan, gfp-ftsZ ($_{352\text{-}AAAAAAA\text{-}358}$)* | This study |
| pEt-$His_6$-Gfp-FtsZ mut-2 | *kan, gfp-ftsZ (ΔC9)* | This study |
| pEt-$His_6$-Gfp-FtsZ Δspace | *kan, gfp-ftsZ (deleted for 359-374)* | This study |
| pEt-$His_6$-Gfp-FtsZ swap | *kan, gfp-ftsZ (residues 352-358 swapped with residues 375-383)* | This study |
| pBad-Gfp-FtsZ | *amp, Para::gfp-ftsZ* | [5] |
| pBad-Gfp-IDR | *amp, Para::gfp-IDR* | This study |

**Supporting References**


[1] Blattner FR, Plunkett G, 3rd, Bloch CA, Perna NT, Burland V, Riley M, et al. The complete genome sequence of Escherichia coli K-12. Science. 1997;277:1453-62.
[2] Camberg JL, Hoskins JR, Wickner S. ClpXP protease degrades the cytoskeletal protein, FtsZ, and modulates FtsZ polymer dynamics. Proceedings of the National Academy of Sciences of the United States of America. 2009;106:10614-9.
[3] LaBreck CJ, May S, Viola MG, Conti J, Camberg JL. The Protein Chaperone ClpX Targets Native and Non-native Aggregated Substrates for Remodeling, Disassembly, and Degradation with ClpP. Frontiers in molecular biosciences. 2017;4:26.
[4] Viola MG, LaBreck CJ, Conti J, Camberg JL. Proteolysis-Dependent Remodeling of the Tubulin Homolog FtsZ at the Division Septum in Escherichia coli. PloS one. 2017;12:e0170505.
[5] Camberg JL, Hoskins JR, Wickner S. The interplay of ClpXP with the cell division machinery in *Escherichia coli*. J Bacteriol. 2011;193:1911-8.

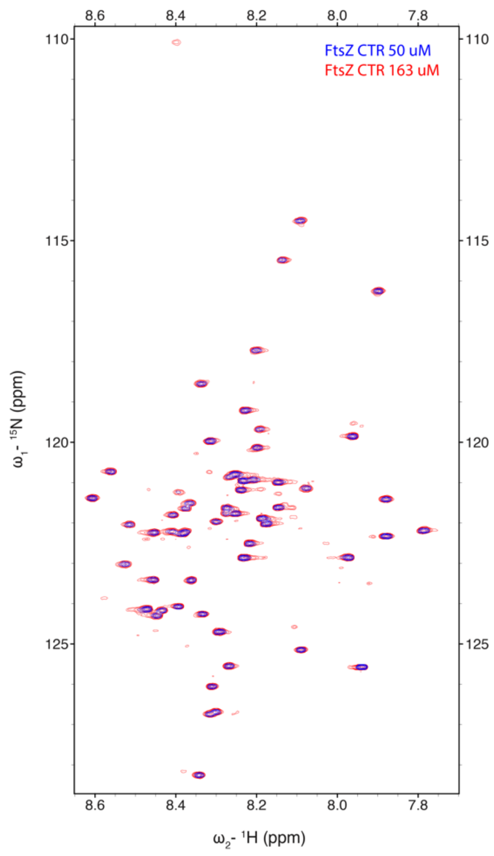


**Fig. S1. The C-terminal region of FtsZ does not self-interact.** $^{1}H$-$^{15}N$ HSQC spectral overlay of 50 μM FtsZ CTR (blue) and 163 μM (red) shows no concentration-dependent chemical shift deviations.

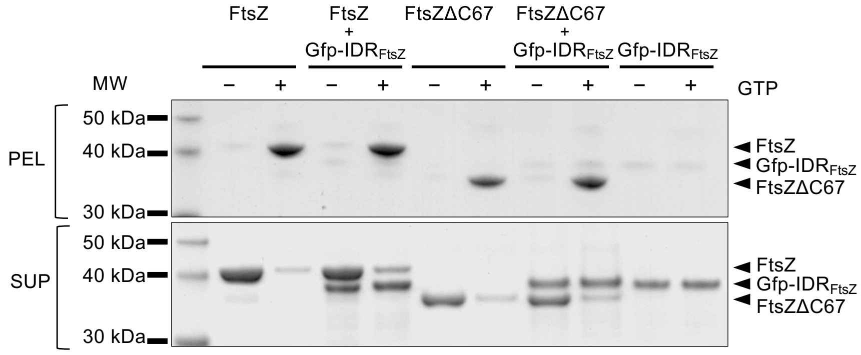


**Fig. S2. FtsZ polymers do not recruit IDRs in sedimentation assays.** Polymers of FtsZ and FtsZ(ΔC67) were assembled with GTP (2 mM) in reactions containing 8 μM FtsZ or FtsZ(ΔC67) in the absence of presence of Gfp-IDR (8 μM). Polymers were collected by ultracentrifugation. Supernatant (SUP) and pellet (PEL) fractions were analyzed by SDS-PAGE and Coomassie staining.
